# Dietary protein restriction diminishes sucrose reward and reduces sucrose-evoked mesolimbic dopamine signaling

**DOI:** 10.1101/2024.06.21.600074

**Authors:** Chih-Ting Wu, Diego Gonzalez Magaña, Jacob Roshgadol, Lin Tian, Karen K. Ryan

## Abstract

**Objective:** A growing literature suggests manipulating dietary protein status decreases sweet consumption in rodents and in humans. Underlying neurocircuit mechanisms have not yet been determined, but previous work points towards hedonic rather than homeostatic pathways. Here we hypothesized that a history of protein restriction reduces sucrose seeking by altering mesolimbic dopamine signaling.

**Methods:** We tested this hypothesis using established behavioral tests of palatability and motivation, including the ‘palatability contrast’ and conditioned place preference (CPP) tests. We used modern optical sensors for measuring real-time nucleus accumbens (NAc) dopamine dynamics during sucrose consumption, via fiber photometry, in male C57/Bl6J mice maintained on low-protein high-carbohydrate (LPHC) or control (CON) diet for ∼5 weeks.

**Results:** A history of protein restriction decreased the consumption of a sucrose ‘dessert’ in sated mice by ∼50% compared to controls [T-test, p< 0.05]. The dopamine release in NAc during sucrose consumption was reduced, also by ∼50%, in LPHC-fed mice compared to CON [T-test, p< 0.01]. Furthermore, LPHC-feeding blocked the sucrose-conditioned place preference we observed in CON-fed mice [paired T-test, p< 0.05], indicating reduced motivation. This was accompanied by a 33% decrease in neuronal activation of the NAc core, as measured by c-Fos immunolabeling from brains collected directly after the CPP test.

**Conclusions:** Despite ongoing efforts to promote healthier dietary habits, adherence to recommendations aimed at reducing the intake of added sugars and processed sweets remains challenging. This study highlights chronic dietary protein restriction as a nutritional intervention that suppresses the motivation for sucrose intake, via blunted sucrose-evoke dopamine release in NAc.

## 1. Introduction

Animals from fruit fly to human naturally love consuming sucrose [1,2] since sucrose not only provides a great resource of energy for survival but also satisfies our sweet tooth. However, as access to food has become easier in recent decades, sucrose overconsumption has emerged as a growing issue. American adults, for instance, have consumed considerably higher dietary added sugar than suggested by the American Heart Association since 2010 (17.0 tsp/day vs. 6-9 tsp/day recommended) [3,4], and sugar consumption in the US was still steadily increasing in the past decade [5]. High dietary sugar consumption is associated with overweight and obesity [6,7], and increases the risk of non-alcoholic fatty liver disease [8], diabetes [9,10], and other cardiometabolic diseases [9–12]. Identifying nutritional and/or behavioral strategies to promote healthier eating habits, by reducing the motivation to consume sugars, may have significant implications for public health.

A growing literature suggests manipulating protein status could influence the desire to eat sucrose and other sweets. For example, using the nutritional geometry framework, Li *et al.* found *Drosophila* fed a low-protein high-carbohydrate diet (LPHC) were less likely to detect and consume dilute sucrose solution(s) compared to control-fed flies [13]. Likewise, mice maintained on LPHC diet, and simultaneously offered *ad libitum* sucrose solution, drank less sucrose compared to controls despite no change in overall calorie intake [14]. Underlying neurocircuit mechanisms have not yet been determined, but in a similar study LPHC-feeding did not affect the consumption of a less-tasty maltodextrin solution [15]—suggesting to us that protein status likely alters sucrose intake via hedonic rather than homeostatic pathways. In agreement with this, human subjects eating low-protein (vs. high protein) diets for 14 days reported decreased explicit and implicit wanting for sweet (vs. savory) foods [16].

Considerable evidence implicates the mesolimbic dopamine system to facilitate rewarding and motivational aspects of eating. Nucleus accumbens (NAc) dopamine signaling is thought to be critical for the associative learning of food reward value [17–19]. Oral sucrose stimulates dopamine release in the NAc and pharmacologic or chemogenetic inhibition of NAc dopamine receptors reduces *ad libitum* sucrose intake [20]. Further, stimulating dopaminergic neurons projecting from the ventral tegmental area to NAc is sufficient to establish a conditioned place preference (CPP) [21] and conversely, pharmacologic inhibition of dopamine receptors blocks the ability of mice to form a CPP for sucrose [19,22]. Lastly, evidence supports that the incentive effects of sucrose and related cues vary according to the internal state of the organism. For example, hedonic reactivity and incentive motivation for food depends on energy status in both mice and humans [23].

The above literature suggested to us that dietary protein status may influence sucrose intake by altering its reward value, via the mesolimbic dopamine system. We tested this hypothesis using established behavioral tests of palatability and motivation, together with modern optical sensors for measuring real-time dopamine dynamics in NAc with fiber photometry.

## 2. Material and Method

### 2.1 Animals

Age-matched male C57Bl/6J mice, 8 to 14 weeks of age, were used in this study. C57BI/6J mice were obtained from The Jackson Laboratory or were bred in-house up to 3 generations removed from the founders. Mice were singly housed on 12 h light/dark cycle in a temperature (20°C to 22°C) and humidity-controlled vivarium with ad libitum access to food and water unless otherwise noted. All animal experiments were approved by the Institutional Care and Use Committees of the University of California, Davis.

### 2.2 Diets

Mice were maintained on standard chow diet (Harlan, catalog #5008, Madison, WI) until the behavioral tests began. Purified, pelleted control diet (CON, D11051801, 18 kcal% protein: 60 kcal% carbohydrate) and low-protein high-carbohydrate diets (LPHC; D11092301, 4 kcal% protein: 74 kcal% carbohydrate), matched for sweetness, were manufactured by Research Diets (New Brunswick, NJ). Full nutritional details can be found in Table 1. Sucrose pellet (#F0023) were from Bio-Serv (Flemington, NJ)

**Table 1:** Diet composition for pelleted diets.

| **Ingredient (g)** | **D11051801** | **D11092301** |
| --- | --- | --- |
| Casein | 200 | 50 |
| L-Cystine | 3 | 0.75 |
| Corn Starch | 375.7 | 485 |
| Maltodextrin 10 | 125 | 150 |
| Sucrose | 107.0777 | 107.0777 |
| Cellulose | 50 | 50 |
| Soybean Oil | 25 | 25 |
| Lard | 75 | 75 |
| Mineral Mix S10022G | 0 | 0 |
| Mineral Mix S10022C | 3.5 | 3.5 |
| Calcium Carbonate | 12.495 | 8.7 |
| Calcium Phosphate, Dibasic | 0 | 5.3 |
| Potassium Citrate, 1 H2O | 2.4773 | 2.4773 |
| Potassium Phosphate, Monobasic | 6.86 | 6.86 |
| Sodium Chloride | 2.59 | 2.59 |
| Vitamin Mix V10037 | 10 | 10 |
| Choline Bitartrate | 2.5 | 2.5 |
| FD&C Yellow Dye #5 | 0.05 | 0 |
| FD&C Red Dye #40 | 0 | 0.05 |
| FD&C Blue Dye #1 | 0 | 0 |
| Kcals/gram | 4.082 | 4.15 |
| Protein (gm%) | 17.88 | 4.569 |
| Carbohydrate (gm%) | 66.72 | 81.437 |
| Fat (gm%) | 9.99 | 10.154 |
| Protein (kcal%) | 17.52 | 4.38 |
| Carbohydrate (kcal%) | 60.46 | 73.6 |
| Fat (kcal%) | 22.02 | 22.02 |
| Calcium, grams | 5.06 | 5.06 |
| Phosphorus, grams | 3.16 | 3.17 |
| Potassium, grams | 3.6 | 3.6 |
| Folate, grams | 2 | 2 |
| Omega 6 FA | 15.10 | 15.10 |
| Omega 3 FA | 3 | 3 |
| Omega 6:3 ratio | 5.03 | 5.03 |

Abbreviation: FD&C, food, drug, and cosmetic.

### 2.3 Palatability contrast test

To determine the effect of protein status on sucrose reward, we used a previously-established “palatability contrast” test (e.g. [24–30]). Sometimes called the “dessert effect” test, this procedure has been used to investigate hedonic aspects of feeding behavior [31]. Briefly the test mimics the common experience that, even after eating an initial meal or pre-load to satiation, individuals may be motivated to further consume a rewarding ‘dessert’.

First, we established protein restricted and control status, by feeding mice either LPHC or CON diet for 4 weeks. Mice continued eating these diets in week 5, while we performed the palatability contrast test. On the night before behavioral testing, we applied a 40% caloric restriction. That is, all mice were provided only 60% of their daily caloric intake. This was to allow motivation for meal feeding and to synchronize food intake during the test. On the test day, mice were fed an initial 2-hour ‘pre-load’ beginning at ZT 7. This consisted of *ad libitum* access to the previously assigned LPHC or CON maintenance diet. Next, beginning at ZT 9, we additionally provided 1-hour *ad libitum* access to sucrose pellets as ‘dessert’ (Fig. 1A). Note: mice were briefly exposed to the sucrose pellets approximately 1 week before behavioral testing, to prevent neophobia during the test.

**Figure 1.**
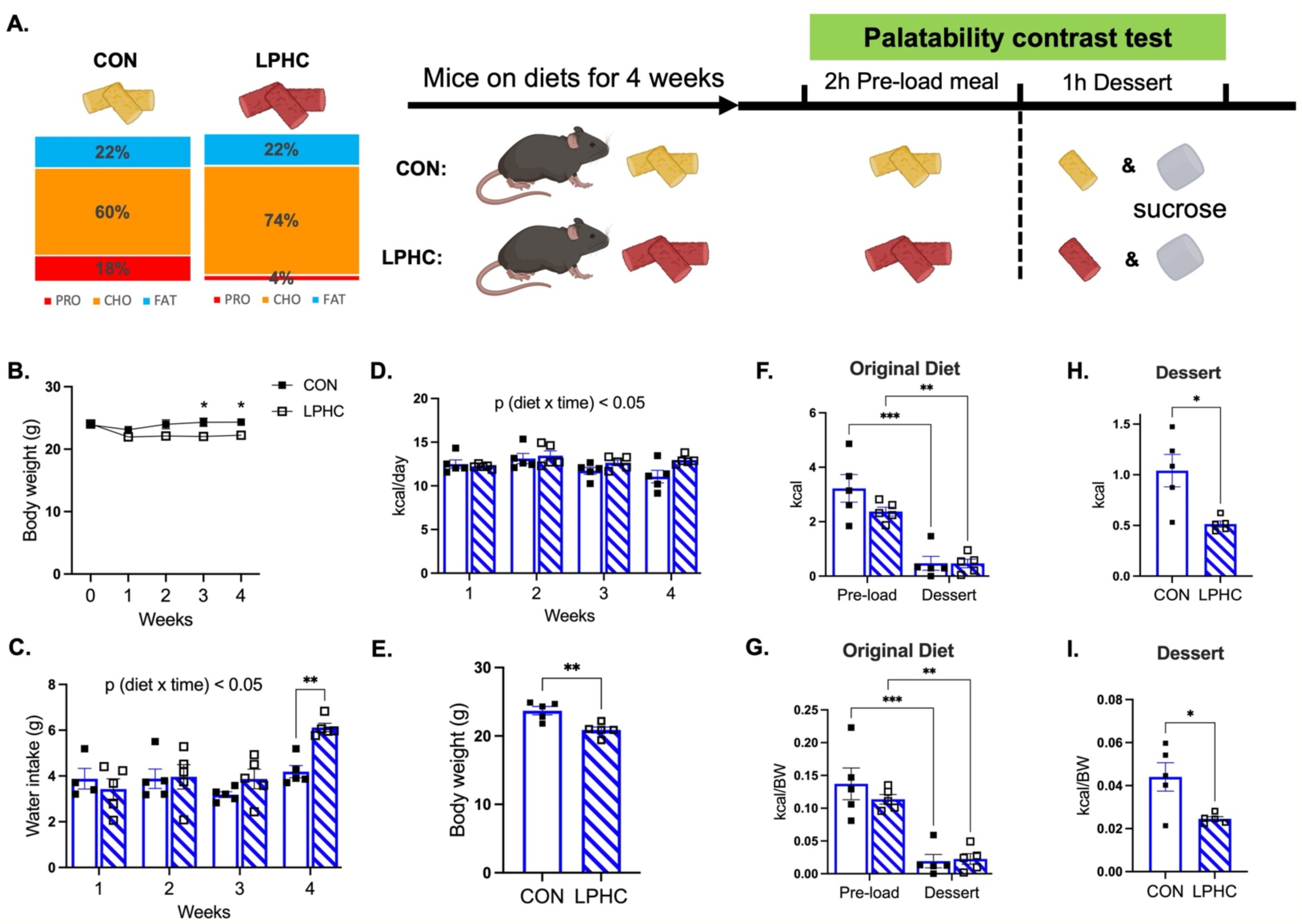
Chronic dietary protein restriction decreased the consumption of sucrose dessert. Mice were maintained on either CON or LPHC diet for 4 weeks before participating in the palatability contrast test, which included a 2h pre-load meal followed by a 1h sucrose ‘dessert’ (**A**). Over the 4 weeks of maintenance diet, LPHC-fed mice lost body weight (**B**) and had greater water intake (**C**) and similar caloric intake (**D**) compared to CON-fed mice. On the day of the palatability contrast test, LPHC-fed mice weighed less than CON-fed mice (**E**). Both CON and LPHC-fed mice consumed the majority of their calories from maintenance diet during the 2h pre-load meal, and ate relatively little maintenance diet during the ‘dessert’ phase of the test (**F, G**). LPHC-fed mice ate significantly less sucrose dessert compared to CON-fed mice (**H, I**). One data point was excluded in week 1 in panel C because of the water bottle leakage. Analyses made by two-way ANOVA with *Sidak post hoc* tests (**C, D, F, G**) or t-test (**B, E, H, I**). Data are shown as mean ± SEM, n= 5 mice per group. *** = p < 0.001, ** = p < 0.01, * = p< 0.05.

### 2.4 Stereotaxic surgery for overexpression of dLight

Anesthetized mice were mounted in the stereotaxic frame (Kopf Instruments), and received subcutaneous injection of anti-inflammatory carprofen (5 mg/kg). An incision was made to expose the skull, and a craniotomy was made above the NAc (AP: +1.3mm, ML: −1.5 or +1.5 mm, DV: −4.25mm). 400 nL AAV9.Syn.dLight virus (2.0 x 10^12^ vg/ml, #240123-01, NeuroTools Vector Core) was slowly injected, unilaterally. An optical fiber (400 µm diameter, 5 mm cut length, NA 0.5, RWD Life Science Inc.) was implanted at 0.05 mm above the virus injection site. A layer of adhesive cement (C&B Metabond, Parkell Inc.) was applied on the skull surface to hold the implanted ferrule, and then a thick layer of resin acrylic (mixture of #0921092 and #0921091, Keystone) was applied to build a head cap. Animals recovered for at least 2 weeks after surgery to allow for dLight expression, and then put on diet treatment as indicated.

### 2.5 Fiber photometry (FP)

We measured sucrose-evoked dopamine release using fiber photometry. Following recovery from the stereotaxic surgery, mice were counterbalanced into diet treatment groups by body weight and maintained on CON or LPHC diets for 5 weeks to establish protein status. Next, we restricted caloric intake for ∼2 weeks, to motivate consumption of sucrose pellets during the test, maintaining body weight at 85-90% of *ad libitum*. On test day, we connected the patch cord, and mice were allowed to freely explore in their home cage while they were recorded from above by the behavior tracking camera (Doric). FP signals and video recording were synchronized following manufacture instructions. We recorded behavior and FP signals in the home cage, while mice consumed 3 sucrose pellets that were sequentially added to the cage. The inter-trial intervals (ITI), between pellets was variable [86.0 ± 53.7 seconds]. To avoid neophobia, all mice had one previous exposure to the sucrose pellets. Likewise, they had been acclimated the patch cord. For behavioral analysis, the onset and completion of each pellet consumption was time stamped, frame by frame, *post hoc*. Both CON and LPHC mice spent ∼20 sec to completely consume each of the 3 sucrose pellets.

Our photometry system employed a 465 nm LED (Doric) for fluorescence excitation and a 405 nm LED (Doric) for isobestic excitation, which were modulated at 211 Hz and 531 Hz, respectively. The sampling rate was 12k Hz. The LED power was in low power mode to acquire stable signals following the manufacturer’s instructions. A fluorescence minicube (Doric) combined both wavelengths, which were then delivered through the optical patch cord to the implanted optic cannula. Emitted light was transmitted through the same patch cord back to the minicube, where filtered for GFP emission wavelength (525 nm) and sent it to the fluorescence detector (Doric). Signals were then demodulated and synchronized with the video recording via the fiber photometry console (Doric). Photometry data were analyzed using custom-written Python scripts. To correct for motion artifact and photobleaching, the 405 nm signal was fit to the 465 nm signal by using the linear squares regression [32]. The fitted 405 nm signal was then subtracted from the 465 nm signals, and then divided by the fitted 405 nm signal to yield ΔF/F values.

Phasic peri-event signals (−10 to +20 sec) were aligned with time stamps representing the onset of the intake for each sucrose pellet. ΔF/F signals were further normalized to z-score ((F_t_ – F_mean_)/F_SD_, where F_mean_ and F_SD_ are the mean and standard deviation from −10∼-5 s prior to the onset of sucrose consumption, as in [33]), to measure the area under the curve (AUC) during the sucrose consumption (0 to +20 sec). Notably, the ΔF/F value from −10∼-5 sec showed no difference between two treatment groups. In addition, tonic dopamine release was revealed by analyzing the mean ΔF/F signals at baseline (before consuming the first sucrose pellet) and during the ITIs. Thus, both baseline and ITI signals excluded 5 sec prior and 20 sec after the sucrose consumption (Fig. 2G).

**Figure 2.**
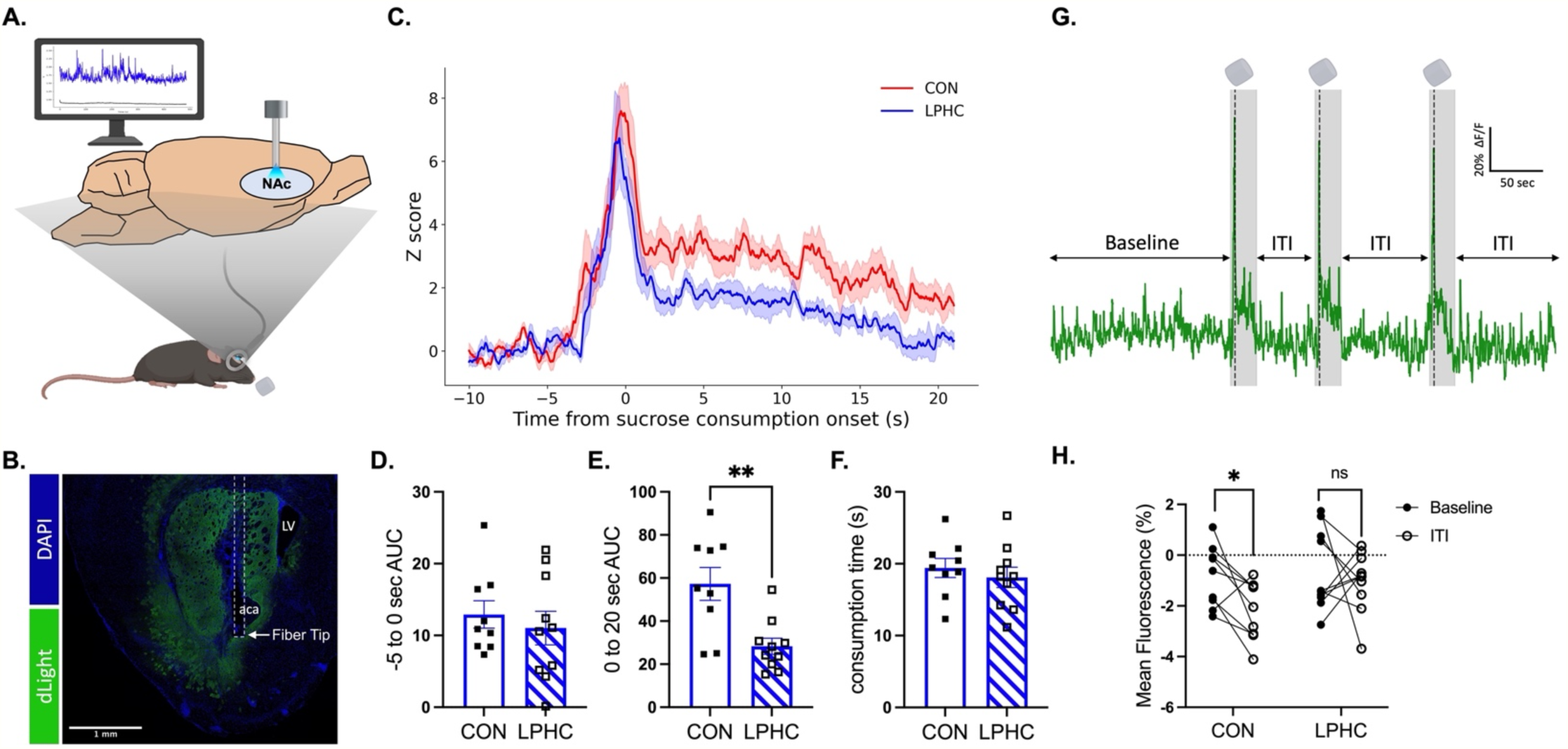
Protein restriction altered sucrose-evoked dopamine dynamics in NAc. An optical fiber was implanted into the NAc, to record dopamine release using dLight fluorescence (**A, B**). Sucrose consumption increased dopamine release in NAc in both CON and LPHC-fed mice, but the magnitude of this effect was significantly blunted with LPHC-feeding (**C**), as measured by the integrated AUC (**E**). There was no effect of protein restriction on dopamine release during the −5 to 0 sec before sucrose consumption (**D**) and no effect on the time spent eating each sucrose pellet (**F**). Sucrose consumption significantly decreased tonic dopamine release in CON-fed mice, as measured at baseline and again during the inter-trial intervals (representative trace shown in panel **G**) and this was abrogated by LPHC-feeding (**H**). Analyses made by t-test (**D-F**) or paired t-test (**H**). Data are shown as mean ± SEM, n= 9-10 mice per group. ** = p< 0.01. LV: Lateral ventricle; aca: anterior commissure; ITI: inter-trial interval. In panel G, dash lines indicate the timing of sucrose consumption, and gray shaded regions indicate 5 sec prior and 20 sec after the sucrose intake.

### 2.6 Conditioned place preference (CPP) apparatus and procedure

To assess motivational aspects of sucrose reward, we used the conditioned place preference test. Behavioral training and testing were conducted in the commercial-available apparatus (Stoeling, catalog #64101, Wood Dale, IL). The CPP apparatus consisted of three chambers; two large conditioning chamber (18 x 20 cm) separated by a shuttle chamber (10 x 20 cm). Conditioning chambers differed from each other in wall patter (circular vs. triangle), and the floor of the entire apparatus was the same. The apparatus was thoroughly cleaned between animals. A GoPro camera was positioned above the apparatus to record animal movement within chambers.

The overall CPP procedure consisted of a pre-test (20 min), six conditioning sessions (30 min), and the place preference test (20 min). To motivate sucrose consumption CON and LPHC-fed mice were calorically restricted to maintain 85-90% of baseline body weight throughout. All procedures were conducted during the light phase (ZT2-10) to allow the use of visual cues. For the pre-test, no food was presented in the chambers. A mouse was placed in the central shuttle chamber and had free access to two conditioning chambers for 20 min. The amount of time the mouse spent in each conditioning chamber was used to calculate a pre-test CPP score (time spent in chamber paired to sucrose minus time spent in chamber paired with empty). Overall, there was no baseline preference for either side, i.e., an unbiased CPP apparatus was used in this study. For conditioning, we assigned the sucrose-paired chamber in a balanced manner, so the mean pre-test CPP score was close to zero for both treatment groups.

Mice were conditioned in the food-paired chamber or unpaired chamber on alternate days, for 6 constitutive days. Each session lasted 30 min. During the sucrose conditioning days, animals consumed ∼5 sucrose pellets in the chamber. On unpaired days, there was no food presented in the chamber in the 30-min session. On the test day, animals were allowed to freely explore all chambers, as the pre-test day with no food present. We used video recording to measure the amount of time in each conditioning chamber.

### 2.7 cFOS Immunolabeling

90 minutes after the CPP test, mice were deeply anesthetized with pentobarbital and transcardially perfused with saline followed by 4% paraformaldehyde (PFA in PBS). Brains were collected and fixed in 4% PFA overnight and transferred to 30% sucrose solution until brains sank. Brains were then sectioned on a freezing microtome (#SM2010R, Leica Biosystems) at 25 µm in a series 1:6, and stored in cryopreservative solution at −20 ℃.

Floating sections were washed (PBS; 3 x 5 min; at room temperature (RT)), blocked (0.3% Triton-X, 5% normal goat serum, PBS; 1h; at RT), and incubated with primary antibody (anti-cFos, #2250, Cell Signaling, 1:1000 in 0.3% Triton-X, PBS; at 4 ℃) overnight. On the next day, sections were washed (PBS; 3 x 5 min; at RT) and incubate with goat-anti-rabbit Alexa Fluor™ 647 secondary antibody (A21244, Invitrogen, 1:500 in 0.3% Triton-X, PBS; at RT). Following another wash (PBS; 3 x 5 min; at RT), samples were incubated with DAPI (D1306, Invitrogen, 1:5000 in PBS; at RT), and mounted on slides.

Sections were imaged by fluorescence microscope (10X: BZ-X series, Keyence; 40X: TCS SP8 STED 3X, Leica), and cFos positive cells were quantified using ImageJ software. Specifically, we counted cells in the NAc core at the level of +1.70 mm to +0.74 mm anterior to bregma, according to The Mouse Brain Atlas [34] using ImageJ. The number of labeled cells were normalized by the area of the outline region of interest across sections. An individual blind to the experimental treatment groups scored the sections. Display images were adjusted for brightness and contrast.

### 2.8 Expression of dopamine receptors and tyrosine hydroxylase

A separate cohort of male mice maintained on CON and LPHC diets for 30 days (from our previous study [35]) were used to examine mRNA expression in the NAc and ventral tegmental area (VTA). Mice were deeply anesthetized with pentobarbital and then decapitated to extract brains, which were immediately frozen in isopentane on dry ice. Coronal sections were cut in 1-mm thick by using a mouse brain-slicing matrix (RBM-2000C, ASI instruments, Warren, MI). The NAc and VTA were identified by referring the architectural landmark [34], and dissected by using 1.50 mm precision brain punch (Ted Pella, Redding, CA) in the cryostat.

Collected punches were homogenized by passage through a 25G syringe. Total RNA was isolated by using a commercially available RNA purification kit (#48500, Norgen Bioteck, Thorold, ON) following the manufacturer’s instructions. cDNA was synthesized using the High-Capacity cDNA Reverse Transcription Kit (ThermoFisher, Cat# 4368814). Quantitative PCR was conducted using Taqman gene expression assay and analyzed on a BioRad CFX384. Expression of dopamine receptor D1 (Drd1; Mm02620146_s1), D2 (Drd2; Mm00438545_m1), and tyrosine hydroxylase (TH; Mm00447557_m1) was normalized to the housekeeping gene 18s ribosomal RNA (Rn18s; Mm04277571) by using the 2 ^-ΔΔCt^ method.

### 2.9 Statistical analyses

Data were analyzed using GraphPad Prism by the appropriate model analysis of variance (ANOVA) or T-test, as indicated. Multiple comparisons were made using *Sidak post hoc* tests. In all cases, α= 0.05. Graphs were created using GraphPad Prism. Data are presented as means +/- SEM unless otherwise noted.

## 3. Results

### 3.1 Low protein status reduced the palatability of sucrose ‘dessert’

LPHC diet-fed mice lost weight despite greater caloric intake [T-test, p < 0.05 (Fig. 1B, D); p(diet x time)<0.05 (Fig. 1B)], and drank more water [p (diet x time) < 0.05, (Fig. 1C)] compared to mice eating CON diet, as expected [35,36]. During the palatability contrast test, both CON and LPHC-fed mice consumed most of their calories from pelleted maintenance diet during the 2-h preload meal and ate relatively little maintenance diet during the ‘dessert’ phase of the experiment [p (time) < 0.001, (Fig. 1F)]. During the 1-h dessert, LPHC-fed mice ate less sucrose compared to controls [T-test, p < 0.05 (Fig. 1H)]. This was true whether the data are considered as an absolute value or were normalized to body weight (Fig. 1G & I). We conclude that chronic dietary protein restriction diminishes sucrose palatability in sated mice.

### 3.2 Low protein status altered both phasic and tonic dopamine release in NAc while mice consumed sucrose pellets in the home cage

Dopamine release in NAc is thought to reflect state-dependent food reward. To determine the influence of protein status on sucrose reward, we used the dopamine sensor dLight, together with fiber photometry, to record sucrose-evoked dopamine release in protein restricted and control-fed mice. First, we overexpressed dLight in NAc, via stereotaxic injection of AAV.Syn.dLight. Following recovery from surgery, mice were maintained on either CON or LPHC diet for 5 weeks. Next, we recorded dopamine release in NAc, during the voluntary consumption of sucrose pellets in the home cage (Fig. 2A, B). We observed a significant increase in fluorescence concordant with onset of eating, as expected [20,33]. Sucrose-evoked dopamine release was observed in both CON and LPHC-fed mice but was significantly reduced in mice with low protein status. That is, when we analyzed peri-event signals by Z-score, LPHC-fed mice had a significantly reduced area under the curve (AUC) measured from 0-20 seconds after the onset of eating [T-test, p<0.01 (Fig. 2C, E)]. We did not observe a difference in the AUC as mice approached the pellets from −5 to 0 sec (Fig. 2D). The average time spent to eat the pellet was similar between the two diet groups (∼20 seconds, Fig. 2F). Moreover, in CON-fed mice, sucrose consumption led to a decrease in tonic dopamine release measured during the inter-trial-intervals (ITI), but this was abrogated by LPHC-feeding [Paired T-test, p<0.05 (Fig. 2G, H)].

### 3.3 Low protein status eliminated the conditioned place preference for sucrose and reduced CPP-associated neuronal activity on the NAc core (NAcC)

Next, we determined the effect of LPHC diet on the motivation to consume sucrose, using the conditioned place preference test (Fig. 3A). Mice showed no preference for either the ‘circle’ or ‘triangle’ conditioning chamber on the pre-test day, regardless of diet treatment, indicating an unbiased apparatus (Fig. 3B). On test day, CON-fed mice exhibited a significant place preference for the sucrose-paired chamber [Paired t test, p<0.05 (Fig. 3C)], as expected [37–42]. This was not observed in LPHC-fed mice [Paired t test, p=0.6983 (Fig. 3C)].

**Figure 3.**
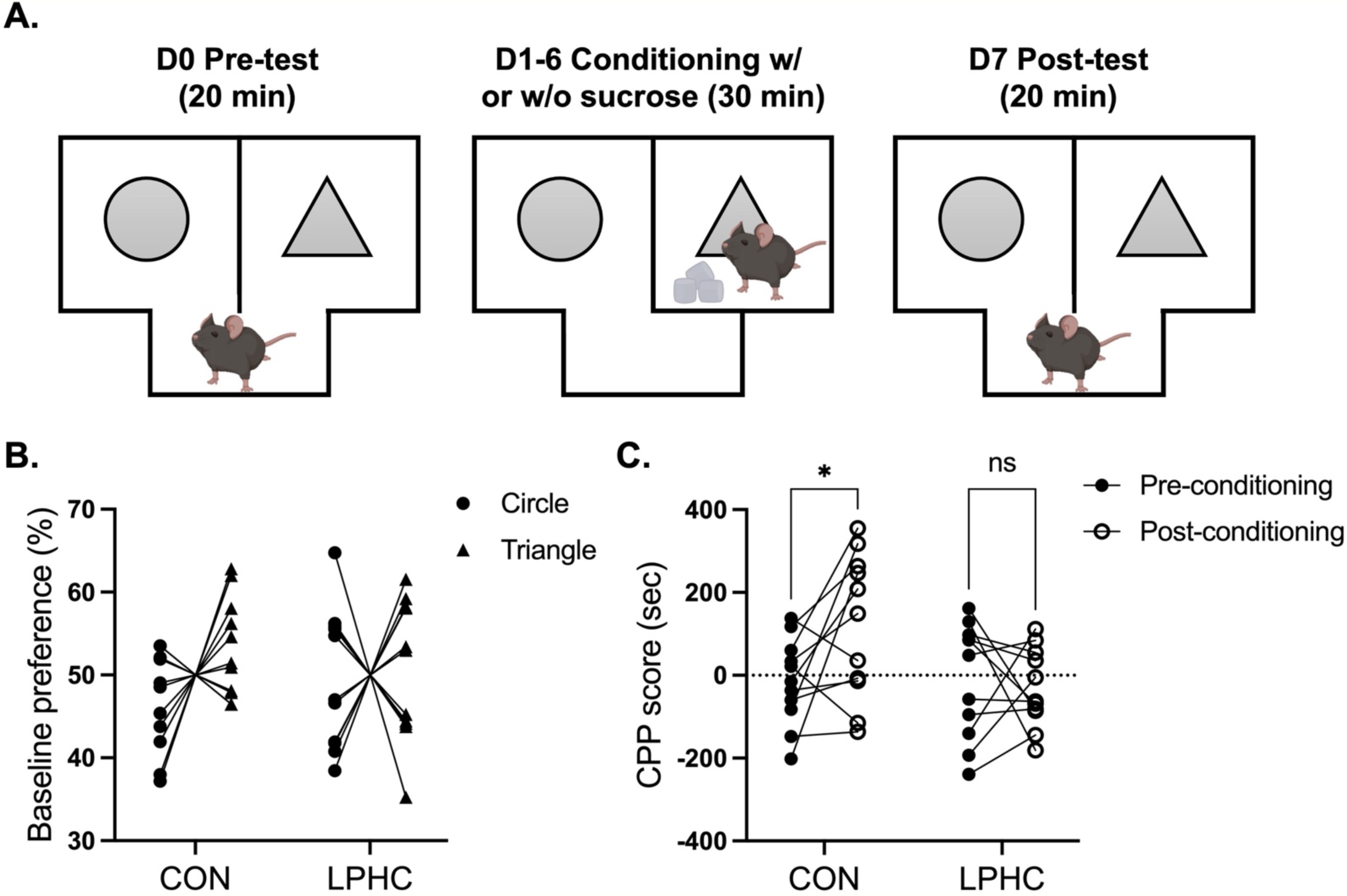
Protein restriction inhibited the sucrose-conditioned place preference. In the conditioned place preference (CPP) test, mice had free access to two conditioning chambers decorated with different wall patterns. Time spent in each chamber was measured before and after 6 days of conditioning (**A**). Prior to conditioning, neither CON or LPHC-fed mice exhibited a preference for the circle vs. triangle-decorated chamber indicating an unbiased apparatus (**B**). After 6 days conditioning with sucrose reward, place preference was measured again in the absence of food cues. CON mice increased time spent in the sucrose-paired chamber; this was not observed in LPHC-fed mice (**C**). Analyses made by paired t-test. Data are shown as mean ± SEM, n= 11 mice per group. * = p< 0.05.

The NAcC is essential for encoding the value of reward-predictive cues and controlling behavioral responses. Accordingly, sucrose seeking behavior increases neuronal activity in the NAcC [17]. To determine whether protein status differently CPP-evoked NAcC neuronal activity, we measured the expression of cFos from mice euthanized 1.5 hours after completing the CPP test (Fig. 4A). We observed fewer cFos positive cells in LPHC-fed mice compared to controls [T-test, p<0.05 (Fig. 4B-D)]. We conclude that low protein status diminishes the motivation to consume sucrose, and this was accompanied by expected differences in the NAcC response.

**Figure 4.**
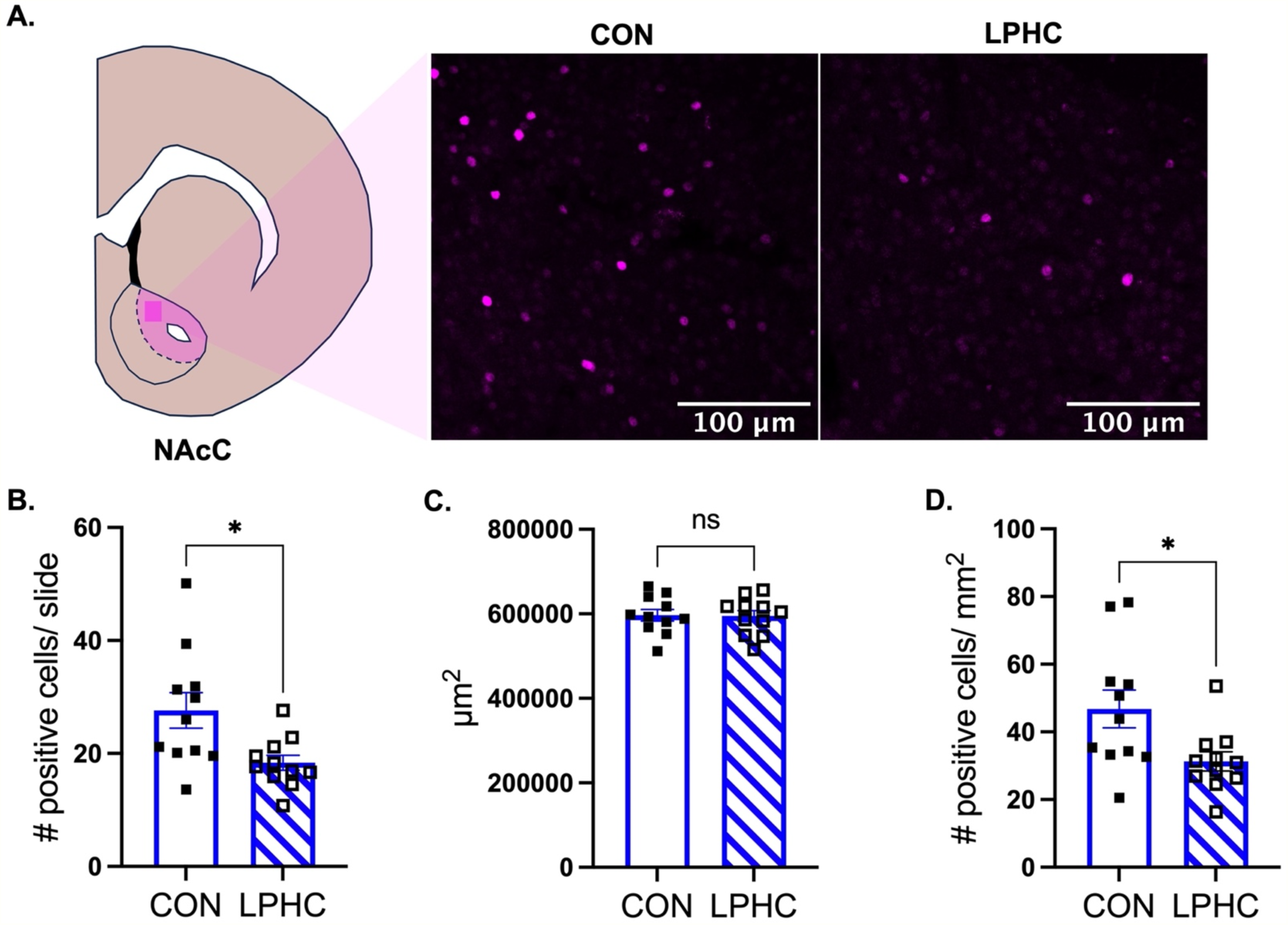
Protein restriction reduced neuronal activity in the NAc core, following the conditioned place preference test. Representative images of cFos immunolableling in the NAcC from CON (A) and LPHC-fed (**B**) mice euthanized 1.5 hours following the CPP test. LPHC-fed mice had significantly fewer cFos-positive cells per slide (**C**) or per mm^2^ (**E**) compared to CON. There was no difference in the area of the NAc used for this analysis (**D**). Analyses were made by t-test. Data are shown as mean ± SEM, n= 11 mice per group. * = p< 0.05.

### 3.4 Chronic LPHC diet did not change mRNA expression for dopamine receptor in the NAc or TH in the VTA

In addition to influencing dopamine release, low-protein status may influence CPP-evoked neuronal activity in the NAc by altering the local expression of dopamine receptors. To explore this, we measured the mRNA expression of DRD1 and DRD2 in the NAc from mice maintained on LPHC and CON diets for 30 days. We found no effect of diet for DRD1/DRD2 expression (Fig. 5A, B). Likewise, there was no effect of protein status on the mRNA expression of tyrosine hydroxylase (TH) in the VTA (Fig. 5C).

**Figure 5.**
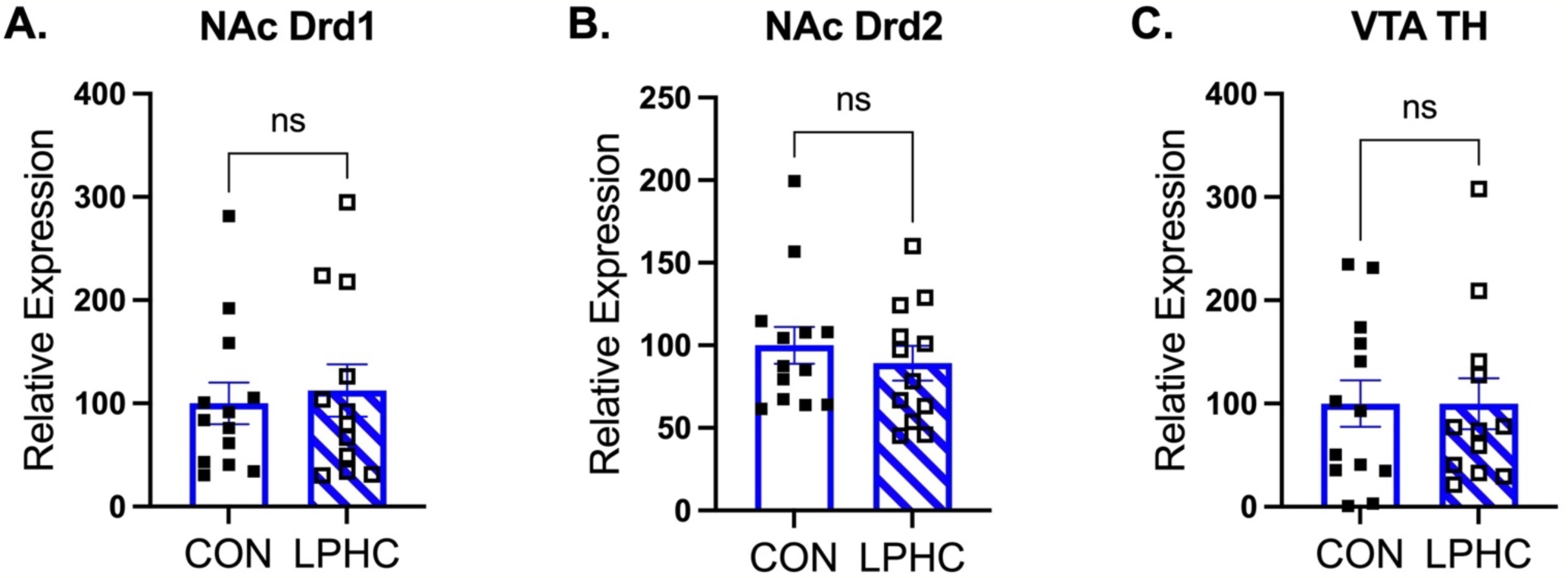
Protein restriction does not alter dopamine receptor mRNA expression in the NAc or TH expression in the VTA. No difference in *Drd1* (**A**) *Drd2* (**B**) or *Th* expression in the VTA (**C**) of mice eating CON vs. LPHC-diet for 30 days. Analyses made by t-test. Data expressed relative to the housekeeping gene, 18s, and shown as mean ± SEM, n= 12-13 mice per group.

## 4. Discussion

Recent findings support that individuals eating a low-protein diet consume less sucrose and other sweets [13,14]. Underlying neurocircuit mechanisms have not yet been determined and may include homeostatic and/or hedonic pathways [43–45]. In this study, we determined the effect of dietary protein restriction on sucrose-evoked hedonic feeding and mesolimbic dopamine signaling in male mice. By identifying the effects of protein status to modulate hedonic hunger (e.g., eating beyond homeostatic need) we provide new mechanistic evidence that suggests protein restriction is a potential intervention for improving overall adherence to a healthy diet.

Using the dopamine reporter dLight and fiber photometry [33], we found that a history of protein restriction reduced sucrose-evoked phasic dopamine release among freely feeding mice in the home cage. Sugar consumption is known to trigger dopamine release in NAc [20] and NAc dopamine is intricately linked to food-seeking behavior, food preference, and the motivation to eat [46–48]. Dopamine neurons are responsive to the nutrient content of ingested food, and this is modulated by prior nutritional status [49–54]. Accordingly, we observed increased dopamine release in NAc of all mice as they consumed sucrose pellets, but this was attenuated in individuals with a history of LPHC-feeding. Low protein status may diminish the rate of catecholamine synthesis in neurons [55–57], by decreasing the availability of the biosynthetic precursors tyrosine and phenylalanine [58–60] and reducing tyrosine hydroxylation [59]. Consistent with this, rats maintained on chronic LPHC diets exhibit decreased dopamine concentration in the striatum, substantia nigra, dentate gyrus [61], and prefrontal cortex [62]. Therefore, it is possible the decrease in sucrose evoked NAc dopamine we observed here simply reflects reduced dopamine availability and a general decrease in motivated behavior. Yet this seems unlikely to us since low protein status *increases* the motivation for dietary protein [63,64]. Accordingly, protein-restricted rodents demonstrate significantly elevated neural activity in the VTA while consuming casein [65] and it *increased* NAc dopamine release in response to electrical stimulation, measured using fast-scan cyclic voltammetry in rat brain slices [46]. Together the data are consistent with a model in which phasic NAc dopamine release in protein-restricted individuals varies according to the stimulus, thereby promoting adaptive changes in food reward and motivation.

We found that history of protein restriction also influenced tonic NAc dopamine signaling during the inter-trial interval, when no stimuli were present. In CON-fed mice, dLight fluorescence dropped below baseline following the consumption of each sucrose pellet; this did not occur in LPHC-fed mice. Tonic dopamine signaling can influence the intensity of a subsequent phasic dopamine response [66,67]. Greater tonic dopamine is thought to inhibit hedonic consumption, e.g. of sucrose [68] or alcohol [69]. Therefore, the differences in ITI dopamine are consistent with the observed effects of protein restriction to inhibit the subsequent dopamine response, and to inhibit sweet ‘dessert’ consumption as we observed in the palatability contrast test.

Building upon evidence that protein context impacts the desire for sweet vs. savory foods in humans [16,70], we found the motivation to consume sucrose was diminished by chronic LPHC feeding in mice using the conditioned place preference test. The CPP is widely used as to measure motivation for drugs of abuse [71–73]. Similarly, sucrose is very effective to condition a place preference in mice [39,42,74]. Here, we observed that control-fed mice exhibited a significant CPP for the sucrose-paired chamber as expected; this was abrogated among LPHC-fed littermates tested in the same experiment. Thus, protein status diminished the incentive motivation for sucrose. Given the association between neuronal activity in the NAcC and cue-induced incentive motivation [50,75–77], we examined c-Fos immunolabeling in the NAcC immediately following the CPP test. In agreement with the behavioral data, LPHC-fed mice displayed significantly fewer c-

Fos-positive cells compared to controls, suggesting a crucial role for neural signaling in the NAc in mediating the LPHC-induced inhibition of sucrose intake. We did not observe changes in the expression of *Drd1*/*Drd2* and *Th* mRNA in the NAc and VTA, respectively, in chronic LPHC-fed mice, despite some studies indicating that protein status may influence the expression of dopamine receptors in the mesolimbic system [61,78]. Overall, chronic LPHC diets did not impact the baseline expression of dopamine receptors and TH but attenuated the motivation for sucrose consumption, accompanied by reduced neuronal activity in NAcC in male mice. Additional work will be needed to identify the specific neuronal populations that were differentially activated during the CPP, but nonetheless these data link diminished motivation for sucrose to blunted NAc activity in protein-restricted mice.

Because sugar and other sweets are rewarding, the overconsumption of sweet foods often occurs even after an individual has just eaten a full meal to satiation. Here we report this so-called ‘dessert effect’ can be modulated by chronic protein status. When mice were fed LPHC diet for 4 weeks, and then fed a pre-load meal to satiation, they ate less of a sweet ‘dessert’ compared to controls. These findings extend previous research showing protein restriction decreases *ad libitum* sucrose and/or sweet preference [14,79,80], by defining the importance of hedonic, non-homeostatic nervous system control. Recent work also highlights that these outcomes are likely context-dependent [14,81,82]. First, the duration of protein restriction is likely important. In our previous study, mice maintained on LPHC diets for only 3 days did *not* reduce the consumption of a sucrose dessert compared to controls [82], suggesting a longer, chronic experience of protein restriction is needed for this effect, and raising the interesting possibility that protein status elicits plasticity in reward circuitry that develops over time. Consistent with this possibility, the decreased preference for sucrose observed in LPHC-fed rats also developed over time [79]. Availability of other macronutrients may also modulate the effect of protein status on sucrose reward and associated mesolimbic dopamine dynamics. LPHC-fed mice have reduced preference for 10% sucrose (vs water) compared to controls, but this is not observed among mice eating low-protein, low-carbohydrate (high-fat) diet [14].

Additional research will be needed to further define the mechanisms linking dietary protein status to sucrose reward, but one likely mediator is fibroblast growth factor 21 (FGF21). FGF21, a hormone released from the liver, is robustly induced by chronic LPHC diets [35,83–86] and appears to influence macronutrient selection in animals and humans [87–89]. Emerging literature indicates that pharmacological administration of FGF21 suppresses sucrose intake by acting on glutamatergic neurons in various regions of the mouse brain, including the ventromedial hypothalamus (VMH) [90] and paraventricular nucleus of the hypothalamus [90,91]. This behavioral modification appears to be translatable to humans, as evidenced by studies showing that subcutaneous injection of BFKB8488A, an agonist antibody targeting FGF21 receptors, reduced subjects’ craving for sweets and carbohydrates after 22 days [92]. Moreover, a negative association was observed between plasma FGF21 levels and cerebral blood flow in the dorsal striatum and other mesolimbic brain regions following sucrose ingestion [93], suggesting that FGF21-induced behavioral modifications may involve the regulation of the mesolimbic system. Consistent with this, recent research demonstrated that an FGF21 analog suppresses alcohol intake by modulating an amygdalo-striatal circuit [94]. Chronic administration of FGF21 also decreased dopamine and dopamine metabolite concentrations in the NAc in mice [95]. Therefore, FGF21 may play a crucial role in regulating sucrose intake in response to chronic LPHC diets by modulating the dopaminergic system. Future experiments are necessary to elucidate the specific role of FGF21 in dopamine signaling and its regulation of sucrose consumption.

Our study has several limitations that warrant consideration. First, because LPHC-feeding influences water intake [36] (and see Fig. 1C), we used sucrose pellets rather than sucrose solution to assess sucrose preference and sucrose-evoked dopamine signaling. We anticipate our findings will generalize to sucrose consumption in various forms, but we acknowledge the need for further research to validate these results. Second, both control and LPHC-fed mice were subjected to caloric restriction to facilitate food consumption in most experiments. It is important to note that caloric restriction may alter dopamine receptor availability [96] and influence patterns of macronutrient selection [97], potentially introducing confounding variables that may affect the interpretation of results. Moreover, there may be cross-interactions between caloric restriction and dietary protein restriction that were not fully explored in this study. Finally, although we observed reduced dopamine release in the NAc during sucrose intake in chronic LPHC-fed mice, the specific upstream neural circuits contributing to this differential regulation remain unclear. The VTA and VMH are potential areas of interest for further investigation. The VTA serves as a major input to the NAc [98], and previous research has demonstrated altered VTA neural activity in protein-restricted rats consuming casein solution [65]. The VMH has been implicated in the regulation of sucrose preference [90], and the hypothalamus may play a role in mediating dopaminergic signaling in the mesolimbic system [48].

## 5. Conclusion

The pervasive issue of sucrose overconsumption, exacerbated by the widespread availability and easy access to sweet ‘junk’ foods, contributes significantly to the global burden of diabetes [9,10], obesity [6,7], and cardiometabolic diseases [9–12]. Despite ongoing efforts to promote healthier dietary habits, adherence to recommendations aimed at reducing the intake of added sugars and processed sweets remains challenging [3–5]. The present study highlights dietary *protein* restriction as a potential nutritional intervention to decrease the motivation for sucrose reward with decreased sucrose-evoked mesolimbic dopamine signaling.

## 6 Acknowledgements

This work was supported by the National Institutes of Health R01DK121035 to KKR. CTW was supported by the Yen Chuang Taiwan Fellowship from the University of California, Davis. Figure 1-3 were created with BioRender.com.

